# Neuroanatomical organization and functional roles of PVN MC4R pathways in physiological and behavioral regulations

**DOI:** 10.1101/2021.03.08.431341

**Authors:** Uday Singh, Kenji Saito, Brandon A. Toth, Jacob E. Dickey, Samuel R. Rodeghiero, Yue Deng, Guorui Deng, Baojian Xue, Zhiyong Zhu, Jingwei Jiang, Leonid V. Zingman, Huxing Cui

## Abstract

**Objective:** The paraventricular nucleus of hypothalamus (PVN) is an integrative center in the brain orchestrating a wide range of physiological and behavioral responses. While the PVN melanocortin 4 receptor (MC4R) signaling (PVN^MC4R+^) is undoubtedly involved in feeding regulation, the neuroanatomical organization of PVN^MC4R+^ pathway and its role in diverse physiological and behavioral regulations have not been fully understood. Here we aimed to better characterize the input-output organization of PVN^MC4R+^ neurons and further test their potential functional roles beyond feeding.

**Methods:** Using a combination of viral tools, we performed a comprehensive mapping of PVN^MC4R+^ circuits and tested the effects of chemogenetic activation of PVN^MC4R+^ neurons on thermogenesis, cardiovascular control and other behavioral regulations beyond feeding.

**Results:** We found that PVN^MC4R+^ neurons broadly innervate many different brain regions known to be important not only for feeding but also for neuroendocrine and autonomic control of thermogenesis and cardiovascular function, including but not limited to preoptic area, median eminence, parabrachial nucleus, locus coeruleus, nucleus of solitary tract, ventrolateral medulla and thoracic spinal cord. Contrary to broad efferent projections, PVN^MC4R+^ neurons receive monosynaptic inputs from limited brain regions, including medial preoptic nucleus, arcuate and dorsomedial hypothalamic nuclei, and supraoptic nucleus. Consistent with broad efferent projections, chemogenetic activation of PVN^MC4R+^ neurons not only suppressed feeding but also led to an apparent increase in heart rate, blood pressure and brown adipose tissue thermogenesis. Strikingly, these physiological changes accompanied an unexpected repetitive bedding-removing behavior followed by hypoactivity and resting-like behavior.

**Conclusions:** Our results clarify the neuroanatomical organization of PVN^MC4R+^ circuits and shed new light on the roles of PVN^MC4R+^ pathways in autonomic control of thermogenesis, cardiovascular function and other behavioral regulations.

## 1. INTRODUCTION

Obesity is a major risk factor for the development of hypertension; population studies indicate that at least two-thirds of the prevalence of hypertension can be directly attributed to obesity[1]. Sympathetic overactivity is well evident in obese individuals and has been considered as one of the major contributors to the development of obesity-associated hypertension[2; 3]. Mounting evidence indicate that the central melanocortin signaling pathway, mainly via melanocortin 4 receptor (MC4R), plays an important role in this pathological process[4-7]. Indeed, in addition to a well-established role in energy homeostasis[8; 9], MC4R signaling pathways are known to affect sympathetic activity and cardiovascular physiology. Central administration of MC4R agonist steadily increases blood pressure (BP) and sympathetic traffics to various organs including the kidney[10-12]. In contrast to elevated sympathetic tone and BP commonly observed in obesity, morbidly obese humans and rodents due to monogenic MC4R-deficiency exhibit low to normal SNA and BP[13-17], suggesting a required but dissociable role of MC4R signaling in both metabolic and cardiovascular regulations in the context of obesity. Notably, a wealth of evidence indicates that MC4R is required for leptin-induced sympathoexcitation; both pharmacological and genetic blockade of MC4R abolish leptin-mediated increase in sympathetic traffic to the kidney[11; 12; 18]. Moreover, central infusion of MC4R antagonist dramatically decreases BP even in spontaneously hypertensive rats with normal body weight[17], proving an important role of MC4R signaling pathway in the neurogenic hypertension beyond obesity. While these publications underscore physiologically important roles for brain MC4R signaling in sympathetic activity and BP regulation, the neural basis underlying MC4R regulation of sympathetic control of cardiovascular function remains incompletely understood.

The paraventricular nucleus of hypothalamus (PVN) is an indispensable integrative center in the brain coordinating a wide range of physiological and behavioral responses, such as feeding, metabolic homeostasis and sympathetic control of cardiovascular function, via its connections to diverse brain regions including different brainstem nuclei and spinal cord[19]. Importantly, the PVN has the highest expression of MC4R among other hypothalamic and brainstem nuclei[20; 21] and PVN^MC4R^ signaling has been well known for its role in the regulation of feeding and body weight homeostasis[22-24]. Additionally, pharmacological study has shown that PVN^MC4R^ signaling may also affect sympathetic activity and BP regulation. Microinjection of MC4R agonist directly into the PVN of anesthetized mice increases BP and sympathetic traffic to the kidney through canonical MC4R-Gαs-cAMP-PKA pathway[25], which is supported by an independent study showing that the ability of systemic MC4R agonist to increase BP is lost in mice lacking Gαs in entire PVN neurons[26]. Of note, slice electrophysiological study has shown that MC4R agonist increases the firing of rostral ventrolateral medulla (RVLM)-projecting pre-sympathetic PVN neurons in Zucker rats[27], with a greater increase of firing activity in obese rats compared to lean control, suggesting an enhanced activity of PVN^MC4R^ signaling pathway may underlie obesity-associated hypertension. While these pharmacological and *ex vivo* electrophysiological approaches support a role for PVN^MC4R^ signaling in sympathetic control of cardiovascular function, the neuroanatomical organization of PVN^MC4R^ pathways and the effects of selective activation of PVN^MC4R+^ neurons on cardiovascular and other behavioral and physiological parameters have not yet been fully examined.

In the present study, we performed a comprehensive neuroanatomical mapping of PVN^MC4R^ circuits using a combination of viral tools and tested the hypothesis that indiscriminate activation of PVN^MC4R^ neurons not only affects feeding but also cardiovascular function and other behavioral physiological parameters through their diffuse projections to brain regions associated with autonomic and behavioral responses.

## 2. METHODS

### 2.1. Animals

MC4R-2a-Cre knock-in mice (Jackson Stock No: 030759) were obtained from Dr. Brad Lowell group[24]. Mice were group housed in University of Iowa’s vivarium in temperature-controlled environment (lights on: 06:00-Light off: 18:00) with free access to standard chow diet and water. Mice used in the present study were all maintained in C57BL/6 and 129 mixed background. All animal protocols were approved by the University of Iowa’s, Institutional Animal Care and Use Committee, and are in accordance with NIH guidelines for the use and care of Laboratory Animals.

### 2.2. Viral vectors for neuronal tracing

Viral vectors from different scientific vendors were used to examine the neuroanatomical organization and the functional roles of PVN^MC4R+^ circuits, including Cre-dependent AAV-DIO-ChR2-eYFP (Addgene), AAV-DIO-hM3Dq-mCherry (Addgene), AAV-retro-DIO-Flp (Duke Viral Vector Core); Flp-dependent AAV-fDIO-TVA-mCherry (Salk Institute Viral Vector Core) and AAV-fDIO-G (Salk Institute Viral Vector Core); and glycoprotein-deleted rabies virus (RV-EnvA-ΔG-GFP, Salk Institute Viral Vector Core).

### 2.3. Stereotaxic surgery

Stereotactic surgery was performed as previously described [28-31]. Briefly, male MC4R-Cre mice were deeply anesthetized by intraperitoneal (IP) injection of ketamine/xylazine (100:10 mg/kg) and placed on a Kopf stereotaxic apparatus (Tujunga, CA). Following standard disinfection procedure, ~1.0 cm incision was made to expose the skull and a small hole was drilled into the skull bilaterally at defined positions to target the PVN (coordinates: AP -0.8 mm, ML +1.1 mm, DV -4.9 mm with 10-degree injection arm). Pulled glass micropipette filled with viral vector was slowly inserted to reach targeted brain region and a small volume (150-200 nl) of injection was made by applying pulse pressure using Tritech pressure Microinjector (Tritech Research, Los Angeles, CA). After 10 minutes of waiting to ensure full penetration of injectant into the targeted area, the needle was slowly removed, and the incision was closed by wound clips. Mice were then kept on a warming pad until awake and fed a regular chow diet throughout the experimental period unless otherwise noted.

### 2.4. Fluorescent double immunohistochemistry (IHC)

Brains of mice that received viral injection were processed for histological verification of correct stereotaxic targeting. IHC was performed as previously reported [28; 31]. Briefly, the mice were transcardially perfused with cold PBS followed by 10% neutralized formalin and the brains were removed and immersed in 25% sucrose solution, cut into five series of 30 µm sections, and then stored in cryoprotectant at -20 °C until processed for IHC. One series of brain sections were rinsed in PBS, blocked in 3% normal donkey serum and 0.3% Triton X-100 in PBS for 30 min at room temperature. Sections were then incubated with primary antibodies against GFP (Aves Labs), mCherry (Clontech), or c-Fos (CalBioChem) for overnight at 4 °C, and then washed and incubated with Cy2- or Cy3-conjugated secondary antibodies (Jackson ImmunoResearch) for fluorescent visualization as per the manufacturer’s instructions.

### 2.5. Neuroanatomical tracing of input-output organization of PVN^MC4R+^ neurons

For comprehensive mapping of efferent projections PVN^MC4R+^ neurons, adult MC4R-Cre+ male mice (12-20 weeks old) received unilateral stereotaxic injection of AAV-DIO-ChR2-eYFP (~150 nl) into the PVN using glass micropipette pressure injection system as described above. After 3-4 weeks of viral injection, mice were transcardially perfused with 10% neutralized formalin and the whole brains were cut into five series of 30 µm sections and then stored in cryoprotectant at -20 °C until further processed for IHC. Brain sections were processed for fluorescent IHC for eYFP (using chicken anti-GFP antibody, Aves Labs) and the sections were then mounted onto gelatin-coated slides for microscopic imaging. Slides were imaged by a slide-scanning microscope (VS120, Olympus) and the images were captured and analyzed using Olyvia software. Only correctly targeted cases (n=3) with minimal contamination to adjacent brain regions were used for optical quantitative evaluation of efferent projections.

For whole-brain mapping of monosynaptic inputs to PVN^MC4R+^ neurons, a mixture of AAV-DIO-flp (Cre-dependent) and Flp-dependent AAV-fDIO-TVA-mCherry and AAV-fDIO-RV-G was unilaterally injected into the PVN of MC4R-Cre mice using glass micropipette pressure injection system as described above. After 3 weeks of stereotaxic surgery, a small volume (~100 nl) of glycoprotein-deleted rabies virus (RV-EnvA-ΔG-GFP) was injected into the same side of PVN where a mixture of AAV was previously infused. One week after the RV-EnvA-ΔG-GFP injection, the mice were transcardially perfused with 10% neutralized formalin and the whole brains were cut into five series of 30 µm sections and then stored in cryoprotectant at -20 °C until further processed for IHC. One series of brain sections from each brain were processed for fluorescent IHC for GFP and the sections were then mounted onto gelatin-coated microscope slides for imaging. Slides were imaged by a slide-scanning microscope (VS120, Olympus) and the images were captured and analyzed using Olyvia software. Only correctly targeted cases with minimal contamination to adjacent brain regions (n=2) were used for the evaluation of neurons sending monosynaptic inputs to PVN^MC4R+^ neurons.

### 2.6. Feeding and glucose homeostasis

Adult (10-16 weeks) MC4R-Cre male mice (n=7 mice) received stereotaxic injection of AAV-DIO-hM3Dq-mCherry into the PVN were subjected to food intake and glucose measurements. A crossover design is used to evaluate the effects of PVN^MC4R+^ neuron activation on feeding and glucoregulation. For feeding, ad lib-fed mice were divided into 2 groups and received intraperitoneal (IP) injection of either saline or small molecule DREADD receptor agonist clozapine-N-oxide (CNO; 2 mg/Kg) right before the onset of dark cycle (6 PM) and food intake was measured at 2- and 4-hour post-injection. CNO was administered at volume of 10X body weight (g). Same experiment was performed again next day but with crossover treatment. For glucose homeostasis, 2-3 hours fasted mice were divided into 2 groups and measured for baseline glucose using small volume of blood drawn from the tail followed by IP injection of either saline or CNO (2 mg/Kg). Blood glucose was again measured at 30-, 60-, and 120-min post-injection. Same measurement was performed again next day but with crossover treatment. After one week of resting, the mice fasted for 5 hours were subjected to glucose tolerance test (GTT; 1g/Kg) with 15-min pretreatment of either saline or CNO (2 mg/Kg). After one week of resting, GTT was performed again but with crossover treatment.

### 2.7. Body surface temperature

Infrared temperature (IR) measurement was done using a high-resolution infrared camera (A655sc Thermal Imager; FLIR Systems, Inc.) as described previously[32]. Again, a crossover design was used to evaluate the effects of PVN^MC4R+^ neuron activation on thermogenesis. Briefly, mice were imaged on a multilane treadmill during 3 min of slow walking (7 m/min and 15° incline) as published previously[33]. Images were captured at 1fps. Ten images were quantified for each time point using FLIR ResearchIR software (version 3.4.13039.1003) and intensity was averaged for group comparison. After the measurement of baseline temperature, the mice received IP injection of either saline or CNO (2 mg/Kg) and then the body surface temperature was monitored again 1-hour post-injection and the change of temperature from the baseline was calculated and compared between the groups.

### 2.8. Radio-telemeter implantation and blood pressure measurements

Adult (10-16 weeks) MC4R-Cre+ male mice that received stereotaxic injection of AAV-DIO-hM3Dq-mCherry into the PVN were subjected to radio-telemetry implantation (PA-C10, Data Science Instruments) for continuous BP and heart rate (HR) monitoring as previously reported[34]. Briefly, animals were deeply anesthetized with ketamine/xylazine (100mg/10mg/kg) and then kept on heating pad to ensure proper maintenance of body temperature. Topical application of eye ointment was given to minimize corneal desiccation. Radio-telemeters were disinfected in actril solution overnight and then kept in sterile saline prior to surgical implantation. Under aseptic surgical conditions, the catheter of the radio-telemeter was inserted into the left carotid artery and tied securely using 6-0 surgical suture. The transmitter was tunneled subcutaneously from the neck until the unit reached the midabdominal region. The neck incision was sutured closed with 4-0 absorbable cat gut and then further sealed with tissue adhesive (Vet-Bond) along the incision line. Warmed sterile saline (500 µl) was given subcutaneously and the animals were kept on heating pad until awake.

After about 2 weeks of recovery from the surgery, mice were subjected to the measurements of arterial pressure and HR in the conscious unrestrained state in their home cages. A crossover design is used to measure the BP and HR in response to systemic treatment of either vehicle (saline) or CNO (2 mg/Kg). Mice were handled and subjected to mock intraperitoneal (IP) injection daily (at random times) for 4-5 days to minimize the confounding effects of handling and stress induced by IP injection. On the day of testing, mice were given IP injection of either saline or CNO (2mg/Kg) in the morning (~8 AM) and then in the evening right before dark cycle (6 PM) to evaluate circadian effects of DREADD activation of PVN^MC4R+^ neurons on cardiovascular parameters. Same experiment was performed next day with crossover treatment. BP was continuously monitored and analyzed by PhysioTel CONNECT system and LabChart software (ADInstrument). At the end of experiment, mice were transcardially perfused with 10% neutralized formalin and brains were extracted for histological verification of correct targeting of hM3Dq-mCherry in the PVN. Only correctly targeted cases (n=3) were used for final data analysis.

### 2.9. Behavioral measurements in PhenoTyper cages

Behavior characterization was performed using Model 3000 Noldus PhenoTyper chambers (Noldus, Wageningen, Netherlands) as described previously[35]. PhenoTyper chamber has an infrared CCD camera installed on the top, which allows automated tracking of mouse movement throughout the experimental period. White shredded paper bedding was given on the floor to mimic home cage-like environment. The arena floor (30 x 30 cm) was further divided to 4 sub-arenas representing wheel running, food, water, and shelter zones, respectively (Figure 6A). Video tracking and quantification of mouse behaviors were performed using EthoVision XT 15 software (Noldus Information Technology). Mice were acclimated in the chambers overnight prior to the testing. A crossover design was used to evaluate behavioral changes in response to chemogenetic activation of PVN^MC4R+^ neurons. The distance traveled within arena, body mobility, the time spent in food and shelter zones, the number of water licks, and wheel-running activity were quantified and compared between the groups. At the end of experiment, mice were transcardially perfused with 10% neutralized formalin and brains were extracted for histological verification of stereotaxic targeting.

### 2.10. Repetitive bedding-removing behavior

Customized plexiglass round chambers were used for repetitive bedding-removing behavior. This customized chamber has two small holes on the wall as shown in Figure 7A; blue colored hole was used for providing water and red colored hole was left open during the testing. Regular chow diet was provided through a customized stainless food hopper hanged inside wall of the chamber. Mice were singly housed and acclimated overnight in the chamber with shredded white paper bedding before the testing. On the day of testing, the mice were given IP injection of either saline (control) or CNO and kept in the chambers. The mice were then camera recorded for up to 20 minutes to observe behavioral alterations.

### 2.11. Marble burying test

Marble-burying test was conducted as described previously[36] with minor modifications. Two separate groups of control and DREADD mice were tested in the standard mouse housing cages. Test cage was prepared using 5 cm of corn woodchip-based animal bedding material. Twelve glass marbles were evenly placed on the bedding for testing. Both control and DREADD mice were given IP injection of CNO (2 mg/kg) 15 min prior to introducing to the testing cage. The mice were allowed exploring in test cage for 30 min to assess marble-burying behavior. At the end of experiment, marbles buried at least two-thirds under the bedding material were counted and compared between the groups.

### 2.12. Statistical Analyses

Statistical analyses were performed using GraphPad Prism (GraphPad Software, La Jolla, CA) software. Comparisons between saline- and CNO-treated groups were made by Student’s t-test and one-way ANOVA or repeated measure two-way ANOVA with Tukey post hoc analysis as needed. P < 0.05 was considered statistically significant. Data are presented as mean ± SEM.

## 3. RESULTS

### 3.1. PVN^MC4R+^ neurons broadly innervate brain regions involved in feeding, neuroendocrine regulation and autonomic control

In order to understand the neuroanatomical organization of PVN^MC4R^ pathways, we performed viral-mediated anterograde tract-tracing of PVN^MC4R+^ neurons. Unilateral microinjection of Cre-dependent AAV-DIO-ChR2-eYFP was made into the PVN of MC4R-Cre mice. After 3-4 weeks of viral injection, mice were transcardially perfused with 10% formalin and brains were processed for fluorescent IHC. Imaging of eYFP immunoreactivity from precisely targeted cases (n=3 mice; Figure 1A and 1B) revealed that PVN^MC4R+^ neurons broadly innervate different brain regions involved in feeding, thermogenesis, neuroendocrine control and autonomic regulation, including but not limited to bed nucleus of the stria terminalis (BNST), preoptic area (POA), lateral hypothalamic area (LHA), dorsomedial nucleus of hypothalamus (DMH), ventromedial nucleus of hypothalamus (VMH), arcuate nucleus of hypothalamus (ARC), median eminence (ME), paraventricular thalamic nucleus (PVT), periaqueductal gray (PAG), parabrachial nucleus (PBN), locus coeruleus (LC), nucleus of solitary tract (NTS), dorsal motor nucleus of Vagus (DMV), rostral and caudal ventrolateral medulla (RVLM and CVLM), and thoracic spinal cord (TSC) (Figure 1C-Y). Optical evaluation of eYFP fiber density from scanned images is summarized in Supplemental table 1. Notably, the projections to most brain regions were largely unilateral (Figure S1A-N), except PVA, NTS and DMV where we observed a strong bilateral innervation of PVN^MC4R+^ neurons (Figure 1V and Figure S1C and N).

**Figure 1.**
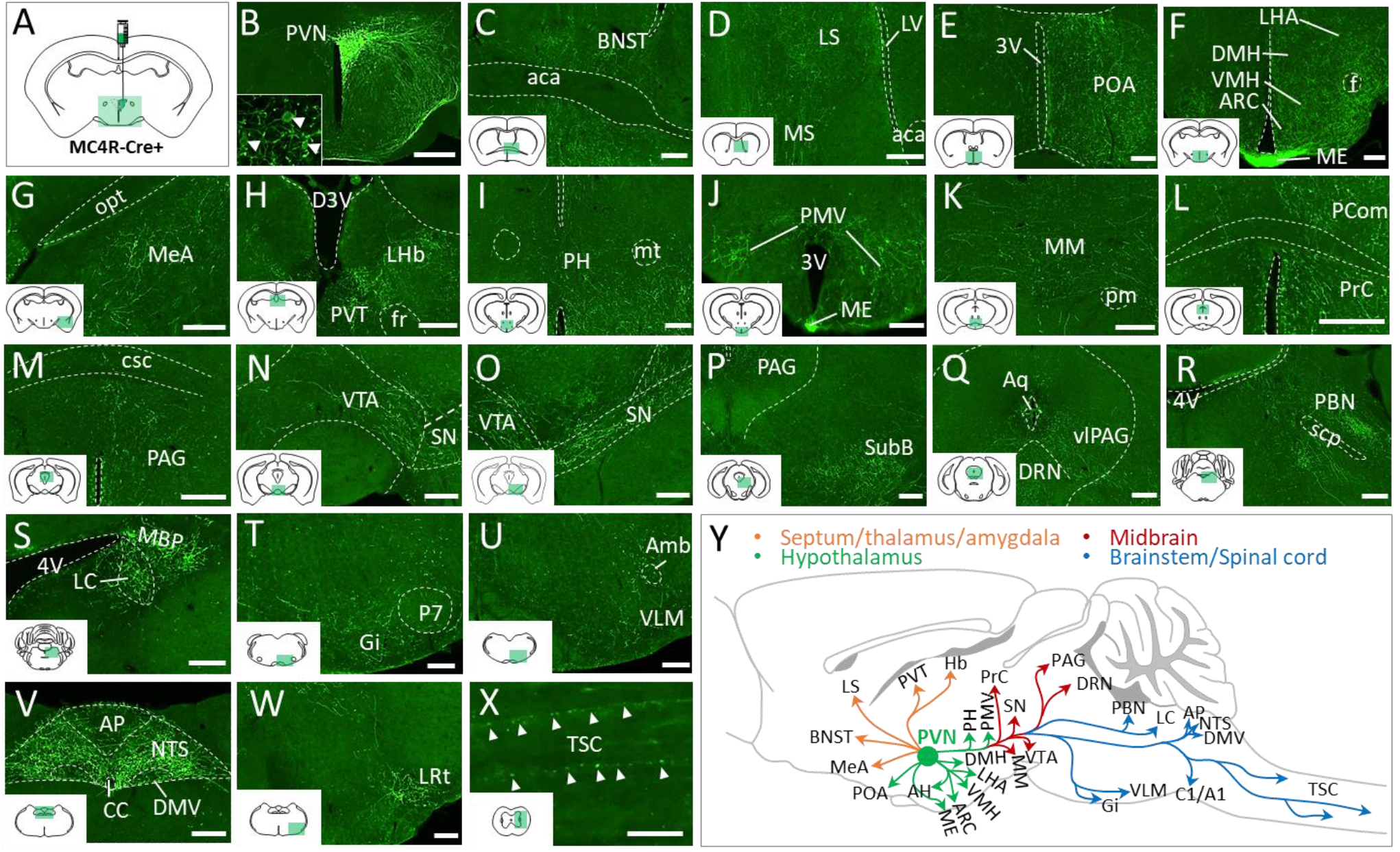
Whole-brain mapping of efferent projections of PVN^MC4R+^ neurons. (A) Schematic showing unilateral viral injection into the PVN of MC4R-Cre^+^ mouse for anterograde tract-tracing. (B) Representative image showing unilateral GFP expression in the PVN^MC4R+^ neurons. Inset shows cell bodies of PVN^MC4R+^ neurons expressing GFP. (C-X) Representative images showing the projections of PVN^MC4R+^ neurons to (C) bed nucleus of the stria terminalis (BNST), (D) lateral (LS) and medial (MS) septal nuclei, (E) preoptic area (POA), (F) dorsomedial hypothalamic nucleus (DMH), ventromedial hypothalamic nucleus (VMH), lateral hypothalamic area (LHA), arcuate hypothalamic nucleus (ARC), and median eminence (ME), (G) medial amygdaloid nucleus (MeA), (H) lateral habenular nucleus (LHb) and paraventricular nucleus of the thalamus (PVT), (I) posterior hypothalamic nucleus (PH), (J) ventral premammillary nucleus (PMV), (K) medial mammillary nucleus (MM), (L) nucleus of the posterior commissure (Pcom) and precommissural nucleus (Prc), (M) periaqueductal grey (PAG), (N) ventral tegmental area (VTA), (O) substantia nigra (SN), (P) subbrachial nucleus (SubB), (Q) dorsal raphe nucleus (DRN) and ventrolateral PAG (vlPAG), (R) parabrachial nucleus (PBN), (S) locus coeruleus (LC) and medial PBN (MBP), (T) Gigantocellular reticular nucleus (Gi), (U) ventrolateral medulla (VLM), nucleus of solitary tract (NTS), dorsal motor nucleus of the vagus (DMV), and area postrema (AP), lateral reticular nucleus (LRt), and (X) few elongated fibers traveling down the thoracic spinal cord (TSC). (Y) Schematic depicting PVN^MC4R+^ axonal projections to the different brain nuclei. Other abbreviations: lateral ventricle (LV), anterior commissure, anterior part (aca), 3rd ventricle (3V), fornix (f), optic tract (opt), dorsal 3rd ventricle (D3V), (fr), mammillothalamic tract (mt), principal mammillary tract (pm), posterior commissure (pc), commissure of the superior colliculus (csc), aqueduct (Aq), 4th ventricle (4V), superior cerebellar peduncle (scp), perifacial zone (P7), ambiguus nucleus (Amb), central canal (CC). Scale bar: 200 µm for B-W, 50 µm for X.

### 3.2. Mapping of monosynaptic inputs to PVN^MC4R+^ neurons

After the whole-brain mapping of afferent projections of PVN^MC4R+^ neurons, we also sought to determine the brain regions where neurons send monosynaptic inputs to PVN^MC4R+^ neurons. To this end, we injected a combination of AAVs driving the expression of TVA and glycoprotein (G) that are required for cell-type-specific monosynaptic inputs into the PVN (Figure 2A-C). After 3 weeks, EnvA-pseudotyped G-deleted Rabies viruses (RV-EnVA-ΔG-GFP) were again injected into the PVN of MC4R-Cre^+^ mouse to label neurons sending monosynaptic inputs to PVN^MC4R+^ neurons as previously reported[37]. In addition to the expected ARC, we observed GFP+ cells in other limited brain regions, including medial preoptic nucleus (MPO), BNST, supraoptic nucleus (SON), ventral subiculum (vSUB), DMH, and NTS (Figure 2D-J).

**Figure 2.**
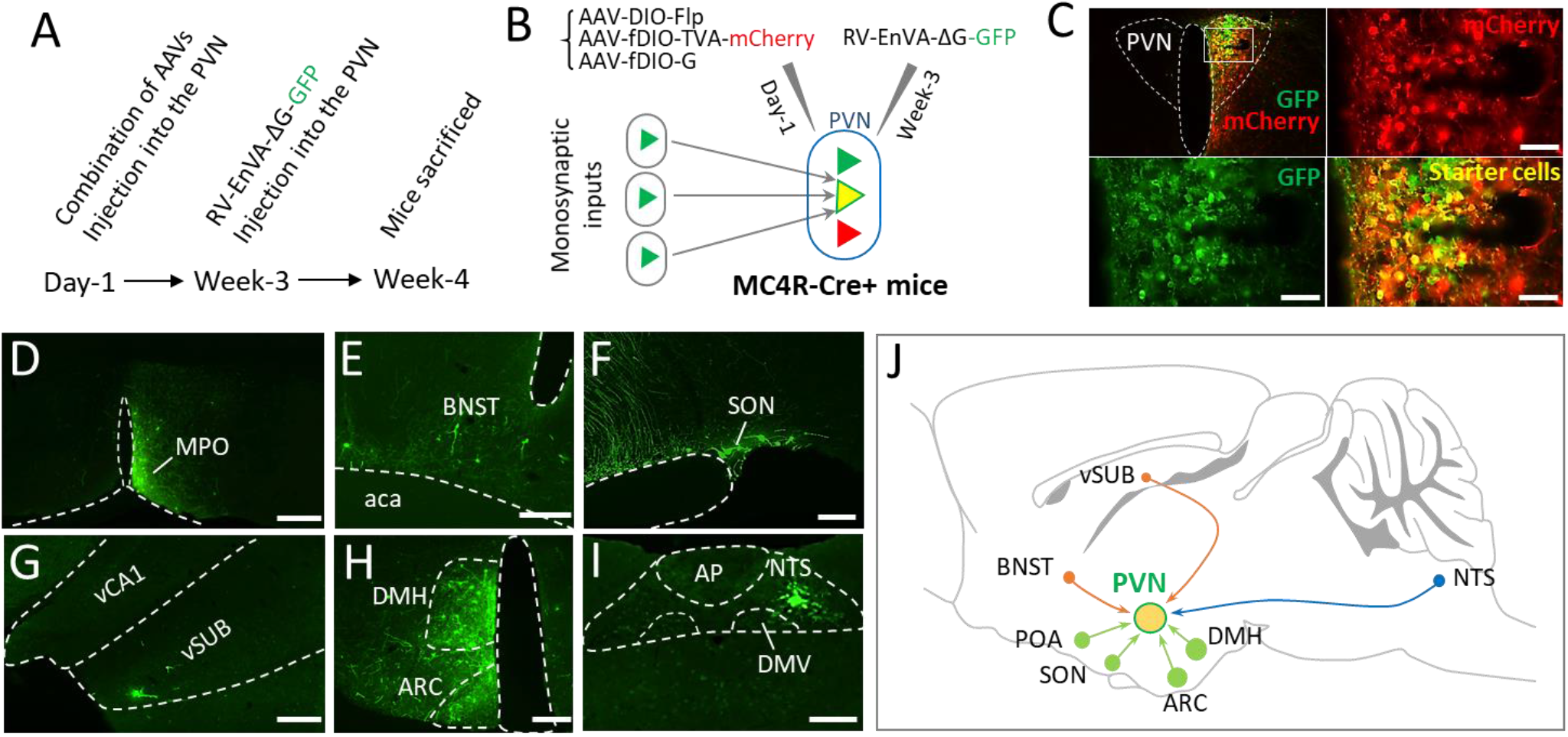
Mapping of monosynaptic inputs to PVN^MC4R+^ neurons. (A) Experimental timeline showing sequential viral injection. (B) Schematic showing the viral strategy to map the neurons sending monosynaptic inputs to PVN^MC4R+^ neurons. (C) Representative images showing the successful targeting of PVN with different viral vectors. (D-I), Representative images showing GFP-positive cells observed in mPOA (D), BNST (E), SON (F), ventral subiculum (vSUB) (G), ARC and DMH (H), and NTS (I). (J) Schematic depicting brain regions where neurons send monosynaptic inputs to PVN^MC4R+^ neurons. Note that the size of the filled circle in each brain region represent relative optical density of observed GFP^+^ cells. Other abbreviations: ventral hippocampal CA1 (vCA1), anterior commissure, anterior part (aca), area postrema (AP), dorsal motor nucleus of the vagus (DMV). Scale bar: 50 µm for C, 100 µm for F, 200 µm for D, E, G, H, and I.

### 3.3. DREADD activation of PVN^MC4R+^ neurons suppress feeding but not glucose homeostasis

To evaluate the functional role of PVN^MC4R+^ neurons, we performed hM3Dq-DREADD-mediated activation of PVN^MC4R+^ neurons. Bilateral microinjection of Cre-dependent AAV-DIO-hM3Dq-mCherry into the PVN of MC4R-Cre+ mice resulted in robust activation of PVN^MC4R+^ neurons as assessed by double fluorescent IHC for mCherry and c-Fos (Figure 3A, B), an established marker of neuronal activation. After confirming effective DREADD-mediated activation of PVN^MC4R+^ neurons, we first tested the effects of DREADD activation of PVN^MC4R+^ neurons on feeding and glucoregulation. As expected, DREADD activation of PVN^MC4R+^ neurons right before the onset of dark cycle significantly suppressed feeding (Figure 3C), which was observed when CNO is administered to control mice without hM3Dq-DREADD expression (Supplemental figure 2A). In contrast, neither baseline glucose nor glucose tolerance was affected by DREADD activation of PVN^MC4R+^ neurons (Figure 3D-E).

**Figure 3.**
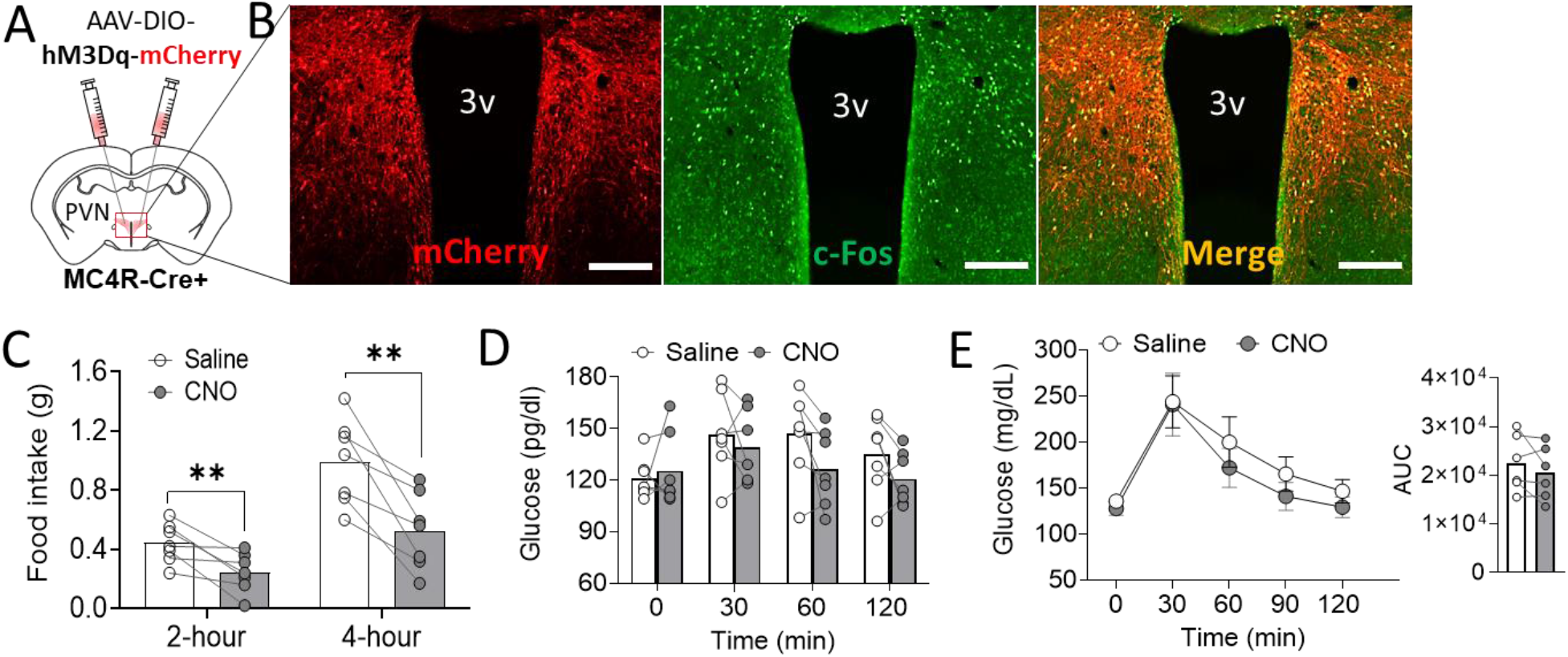
DREADD activation of PVN^MC4R+^ neurons suppress feeding but does not affect glucose homeostasis. (A) Schematic showing bilateral microinjection of Cre-dependent AAV driving expression of excitatory hM3Dq-DREADD receptor into the PVN of MC4R-Cre^+^ mouse. (B) Representative images showing mCherry and c-Fos immunoreactivity in the PVN of MC4R-Cre^+^ mouse after 90 min of CNO treatment (Scale bar: 200 µm). (C) Decrease in food intake following CNO injection at the beginning of dark cycle (6 PM). (D, E) The effects of DREADD activation of PVN^MC4R+^ neurons on fed glucose (D) and glucose tolerance (E). **p<0.01 by paired Student’s t-test (one-tailed). Data are presented as mean ± SEM.

### 3.4. DREADD activation of PVN^MC4R+^ neurons increase thermogenesis

In contrast to the well-established role in feeding regulation, the role of PVN^MC4R+^ pathways in autonomic activity-driven thermoregulation has been vague. Based on an observation of broad innervation of PVN^MC4R+^ neurons to many brain regions involved in sympathetic activation and thermoregulation (Figure 1Y), we next tested whether PVN^MC4R+^ circuits also affect autonomic thermoregulation. To this end, mice were placed on the treadmill for slow walking and body surface temperature was monitored by a high-resolution IR thermal imaging camera as the baseline. Mice then sequentially received IP injection of saline and CNO (2 mg/kg) with 1-hour interval and body surface temperature was taken 1-hour post-injection to evaluate the changes of body surface temperature. While IP saline did not result in notable changes in temperature as expected, there was an obvious increase in body surface temperature after 1-hour IP CNO (Figure 4A). Quantification revealed that DREADD activation of PVN^MC4R+^ neurons significantly increased the temperature in neck area (brown fat), low back and tail (Figure 4B). The changes of temperature in neck area and low back were not observed when CNO was given to control mice without hM3Dq-DREADD expression, however, there was a slight but significantly increase tail temperature by CNO treatment in control mice (Supplemental figure 2B).

**Figure 4.**
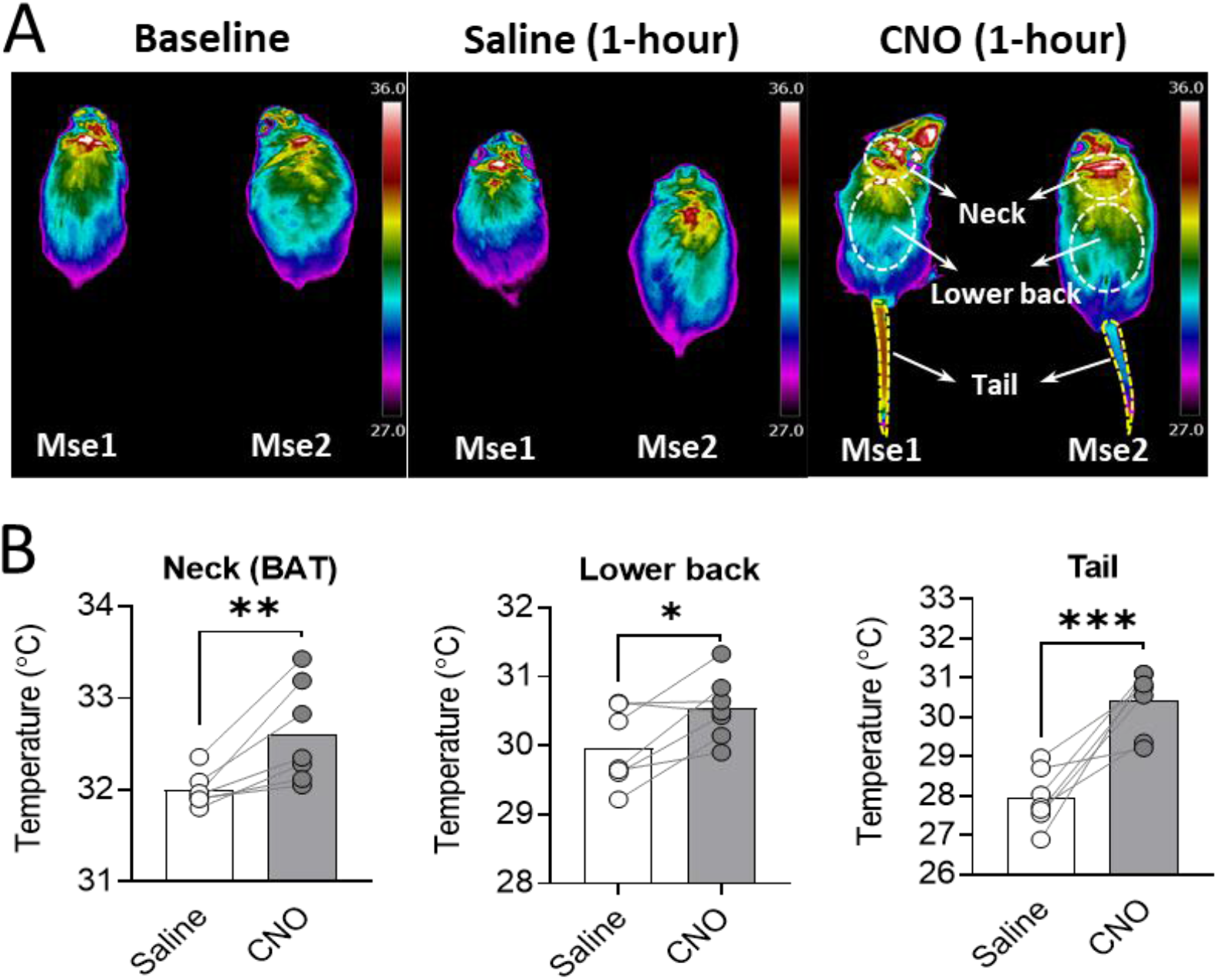
Thermoregulatory effect of DREADD activation of PVN^MC4R+^ neurons. (A) Representative IR thermographic images taken at baseline and after 60 min of saline or CNO treatment while allowing slow walking on a treadmill. Body surface areas of interest were drawn separately for neck (BAT), lower back, and tail. (B) Quantified body surface temperature in mice received saline and CNO treatment (n = 7 mice). *P<0.05, **P<0.01**, ***P<0.01 by paired Student’s t-test.

### 3.5. DREADD activation of PVN^MC4R+^ neurons increase BP and HR

Neurons in the PVN are critical in determining the sympathetic tone to the cardiovascular system and thereby affect BP and HR. Indeed, we observed dense projections of PVN^MC4R+^ neurons to the brain regions that are important for cardiovascular control, including PAG, NTS, RVLM, and TSC (Figure 1). Therefore, we tested whether PVN^MC4R+^ neuron activation also alters BP and HR. To this end, the mice were implanted with radio-telemeter for continuous monitoring of BP and HR. After 2 weeks of recovery from the surgery, mice were tested for their cardiovascular response to IP injection of either saline or CNO in both light and dark phases. Consistent with broad projections to various cardiovascular brain regions, DREADD activation of PVN^MC4R+^ neurons by single dose of CNO (2 mg/kg) resulted in sharp increase in MAP and HR lasting more than 4 hours (Figure 5A-D). Interestingly, these cardiovascular responses elicited by DREADD activation of PVN^MC4R+^ neurons were apparent in light cycle but not dark cycle. Further analysis revealed that DREADD activation of PVN^MC4R+^ neurons resulted in a more prominent increase in systolic arterial pressure (SAP) than diastolic arterial pressure (DAP) in light cycle (Figure 5E-H). None of these changes were observed when CNO was administered to control mice without hM3Dq-DREADD expression (Supplemental figure 2C-F).

**Figure 5.**
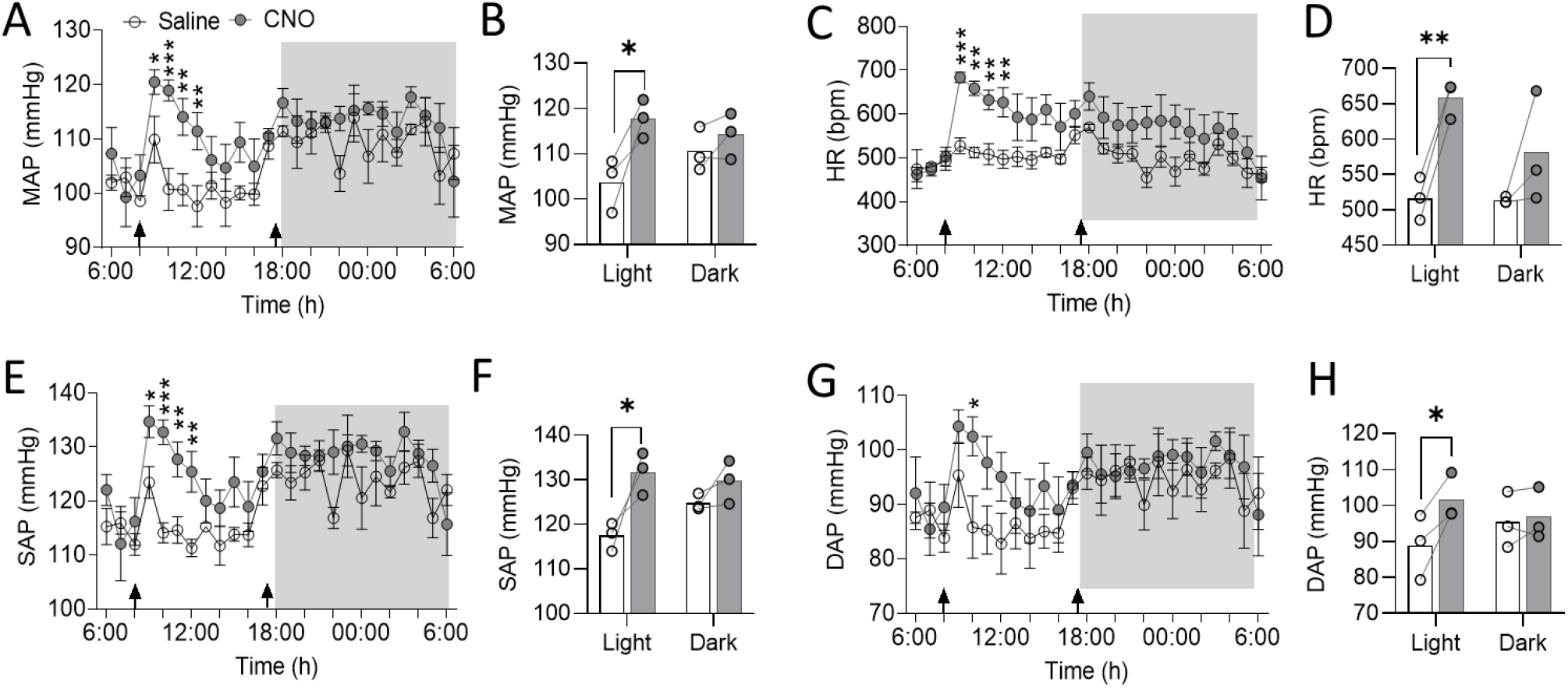
The effects of chemogenetic activation PVN^MC4R+^ neurons on cardiovascular function. MC4R-Cre^+^ mice expressing excitatory hM3Dq-mCherry in the PVN were implanted with DSI radio-telemetry for continuous monitoring of BP and HR. Mice were given IP injection of either saline or CNO in the morning and then again in the evening right before dark cycle (arrows in X-axis indicate the time of each treatment). (A, C, E, G) MAP (A), HR (C), SAP (E), and DAP (G) were analyzed hourly and compared between the groups. (B, D, F, H) The average of 4-hour post-treatment for MAP (B), HR (D), SAP (F), DAP (H) were also analyzed separately for light and dark cycles and compared between the groups. *P ≤ 0.05, **P<0.01, ***P<0.001 by multiple paired t-test. Data are presented as mean ± SEM.

### 3.6. Behavioral alterations by DREADD activation of PVN^MC4R+^neurons

Because of the broad innervation of PVN^MC4R+^ neurons to diverse brain regions that are important for behavioral regulation (BNST, LHA, VTA, PAG, etc), we also tested whether activation of PVN^MC4R+^ neurons elicit other behavioral responses. To this end, we subjected mice to PhenoTyper cages which allow continued monitoring of different mouse behaviors in an automated fashion. After overnight acclimation in the PhenoTyper cages, mice were given IP injection of either saline or CNO right before dark cycle (17:30 – 18:00 PM) and then again next morning (8:00 – 9:00 AM) to determine behavioral alterations in both dark and light cycles. The floor was further marked for 4 sub-arenas representing wheel running, food, water, and shelter zones, respectively (Figure 6A, B), which enable the calculation of time spent in each zone. We found that DREADD activation of PVN^MC4R+^ neurons lead to an initial transient (1-hour post-injection) increase followed by significant suppression of ambulatory activity and body mobility (Figure 6C, D). The mice also spent more time in the shelter zone but less time in the food zone upon DREADD activation of PVN^MC4R+^ neurons (Figure 6E, F). No alteration in water drinking behavior was observed (Figure 6G). Somewhat consistent with reduced ambulatory activity, voluntary wheel running activity was also significantly reduced upon DREADD activation of PVN^MC4R+^ neurons (Figure 6H).

**Figure 6:**
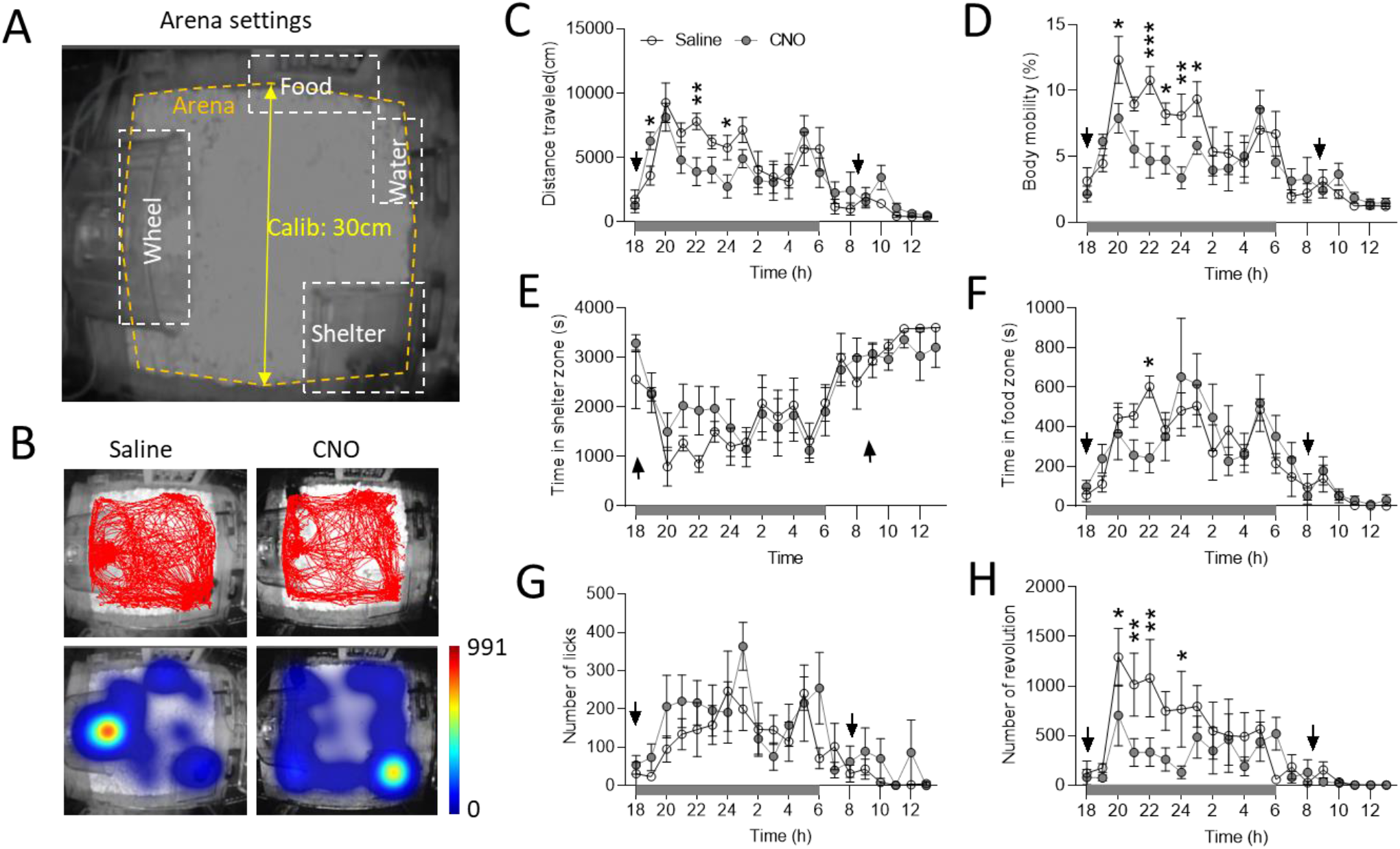
Automated behavioral analysis of mice with DREADD activation of PVN^MC4R+^ in PhenoTyper boxes. (A) Representative schematic showing arena setting in PhenoTyper cage for behavioral characterization. (B) Representative activity trace (upper panel) and heat map (lower panel) showing total distance traveled and the time spent in each zone after either saline or CNO treatment. (C-H) Hourly quantitative analysis showing total distance traveled (C), body mobility (D), time spent in shelter zone (E), time spent in food zone (F), number of water licks (G), and the total number of wheel running (H) in response to saline and CNO treatment (arrows in X-axis indicate the time of injection in the morning and the evening; n = 7 mice). *P< 0.05, **P<0.01, ***P<0.001 by multiple pared t-test. Data are presented as mean ± SEM.

### 3.7. Repetitive bedding-removing behavior by DREADD activation of PVN^MC4R+^neurons

During the measurements of different physiological and behavioral alterations upon PVN^MC4R+^ neuron activation, we noticed a striking repetitive behavior for DREADD mice when tested in their home cages. After ~10 minutes of IP injection of CNO, mice started digging and holding the paper bedding materials and attempted to push them through the wire bar located on the top of cage. To more rigorously monitor this behavior, we tested them in a customized plexiglass chambers while video recording the behavior. After ~10 min of IP injection of CNO, mice started pushing paper bedding out through a hole left open (Figure 7A-C; also see the video of Figure 7D provided in Supplemental Video 1), which was not observed in any of control mice received CNO injection (data not shown). Notably, DREADD activation of PVN^MC4R+^ neurons did not affect marble-burying behavior (Figure 7E, F) – another repetitive behavior implying anxiety or obsessive-compulsive disorder-like behavior.

**Figure 7:**
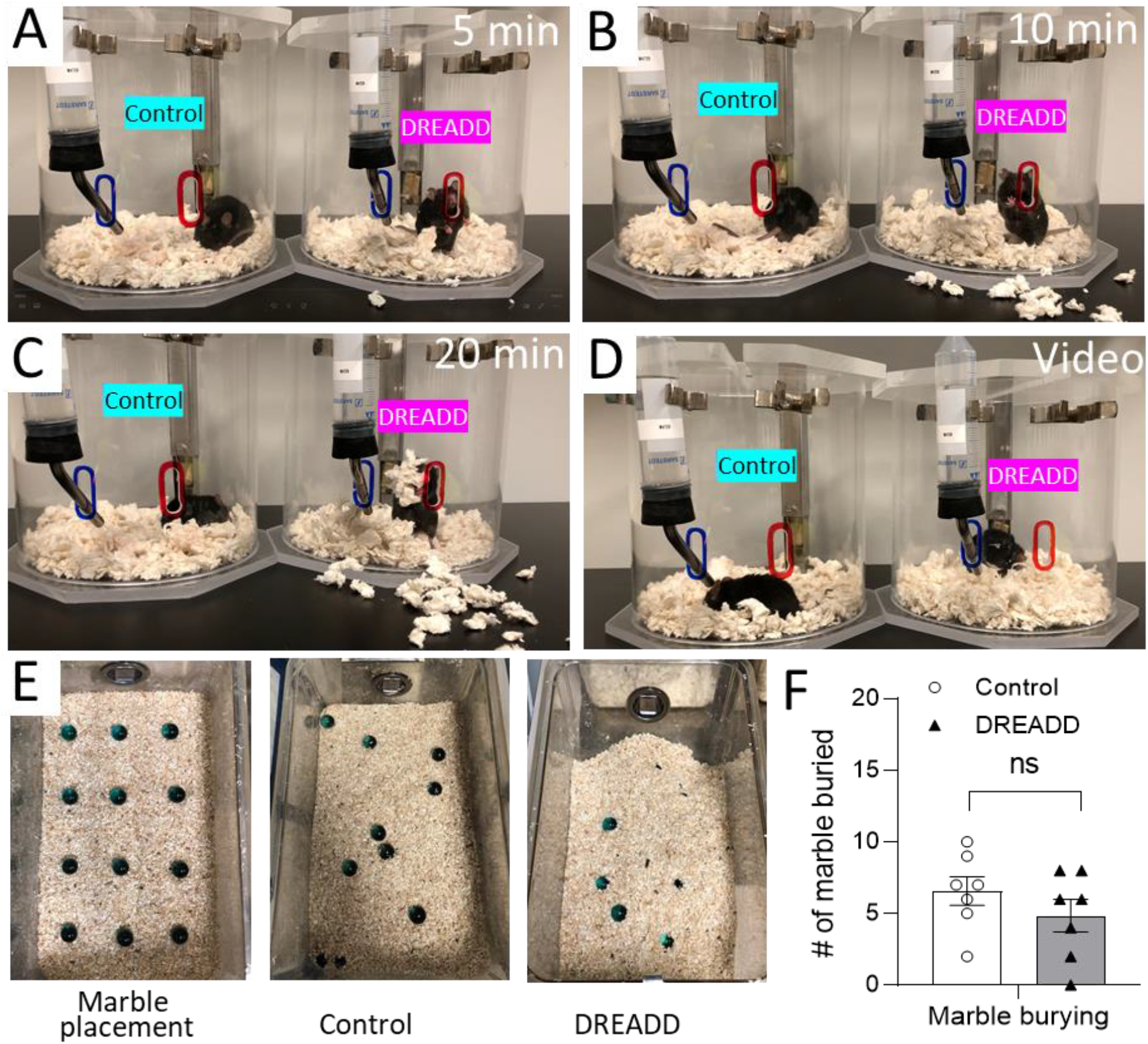
DREADD activation of PVN^MC4R+^ neurons induces repetitive bedding-removing behaviors. (A-C) Representative images showing the removal of paper bedding at 5-min (A), 10-min (B) and 20-min after IP injection of CNO (2 mg/kg). (D) Video demonstration of repetitive bedding-removing behavior following CNO injection (see Supplemental Video 1). (E) Representative images showing the placement of marble before the test and marbles buried upon DREADD activation of PVN^MC4R+^ neurons. (F) Quantification of marbles buried in control and PVN^MC4R+^ DREADD-activated mice by Student’s t-test (n = 7/group). Data are presented as mean ± SEM.

## 4. DISCUSSION

The PVN is a key node in the brain orchestrating a wide range of physiological and behavioral responses, including but not limited to fear, fight-or-flight response, feeding, metabolic homeostasis, and autonomic regulation of cardiovascular function. PVN^MC4R+^ neurons represent a small subset of PVN neurons and are largely distinct from many well-known PVN neurons expressing different neuropeptides[20; 38] and irrefutable evidence support a role for PVN^MC4R+^ pathways in feeding regulation mainly via PVN^MC4R+^→PBN circuit[23; 24; 39]. In the present study, we provide further evidence to show a rather complex role of PVN^MC4R+^ neurons beyond feeding.

We observed a significant increase in body surface temperature around the neck and lower back upon DREADD activation of PVN^MC4R+^ neurons, indicating a likely elevation of sympathetic drive to the BAT. This observation is in line with previous findings that PVN^MC4R+^ neurons polysynaptically innervate the BAT [40; 41], intra-PVN injection of MC4R agonist MTII increase BAT temperature [40], and selective restoration of MC4Rs in Sim-1^+^ neurons that predominantly localized to the PVN rescues blunted MTII-induced increase in energy expenditure in MC4R-null mice [42]. One notable finding, however, from this body surface thermal imaging is a significantly increased tail temperature by DREADD activation of PVN^MC4R+^ neurons. Increased tail temperature is arguably an indication of vasodilation which normally occur in rodents as a mean to promote heat dissipation and reduce body temperature[43]. It is therefore plausible that DREADD activation of PVN^MC4R+^ neurons may also have caused an increase in core body temperature, leading to a counter-regulatory heat-dissipating response by promoting vasodilation. This scenario may also fit to further explain the repetitive bedding-removing behavior which could have been a behavioral thermoregulatory response for mice to promote heat dissipation by eliminating nesting materials. Future experiments measuring the core body temperature and/or cold-seeking behavior in thermogradient environment might help to further prove this possibility. Additionally, an increase in tail temperature (and hence, vasodilation) also indicates that DREADD activation of PVN^MC4R+^ neurons does not seem to cause indiscriminate sympathoexcitation as usually seen in acute stress response, which would have caused vasoconstriction rather than vasodilation. Thus, it is likely that PVN^MC4R+^ neurons selectively influence sympathetic outflows to certain effector organs among others. Another possibility for increased body surface temperature could be due to an evoked neuroendocrine response, such as the activation of hypothalamic-pituitary-thyroid (HPT) axis, which promotes metabolism and heat production. Indeed, most dense innervation of PVN^MC4R+^ neurons was observed in the ME (Figure 1F and Supplemental Table 1). However, since previous studies have shown that there is minimal overlap of PVN^MC4R+^ neurons with thyrotropin-releasing hormone (TRH) [20; 38], it remains to be seen whether PVN^MC4R+^ neurons can activate HPT axis. Functional roles of PVN^MC4R+^→ME/pituitary pathway therefore warrants further investigation.

In addition to metabolic homeostasis, it has been well established that brain MC4R signaling pathways affect sympathetic control of cardiovascular function, which has been an obstacle for developing a safe anti-obesity medication by targeting MC4Rs[6]. MC4Rs are broadly expressed in different brain regions involving in autonomic control of cardiovascular function[20], and our results now support an idea that PVN could be one of the important brain regions where MC4R signaling affects cardiovascular physiology. We observed a sharp increase in MAP and HR upon DREADD activation of PVN^MC4R+^ neurons, especially when neurons are activated in light cycle, but not dark cycle. In fact, the peak responses of BP and HR were even higher when activated in light cycle compared to dark cycle. The underlying mechanism of this light phase-specific rise in BP and HR by PVN^MC4R+^ neuron activation remains unclear but may involves circadian-specific responses of effector organs (vasculature and heart) to a sudden increase in sympathetic outflow. It is worthy to note that single dose of IP CNO in light cycle in fact resulted in a sustained increase in HR and MAP nearly up to 7-8 hours, which might have had an impact on evaluating the effects of the second dose of IP CNO right before dark cycle. Nonetheless, our findings are consistent with previous pharmacological observations showing that intra-PVN injection of MTII increases MAP in anesthetized lean Zucker rats [25] and that MTII-mediated increase in BP is lost in mice lacking Gαs in PVN neurons[26], arguing a plausible role of PVN^MC4R^ signaling in cardiovascular regulation.

Behavioral monitoring in PhenoTyper cages unexpectedly revealed that DREADD activation of PVN^MC4R+^ neurons acutely increase ambulatory movements followed by an obvious behavioral suppression, including reduced locomotor activity, feeding, and voluntary wheel running. These behavioral suppressions indicate that increase in BP and HR is unlikely secondary to the behavioral alterations, implying there could be segregated PVN^MC4R+^ pathways that differentially mediate behavioral and autonomic responses. Indeed, our anterograde tract-tracing revealed that PVN^MC4R+^ neurons broadly innervate many different brain regions known to mediate different behavioral and physiological responses. Because of the well-known cardiovascular side effects of pharmacological activation of MC4R signaling, it is of great interest for future studies to test whether there are anatomically distinct subsets of PVN^MC4R+^ neurons that differentially regulate feeding behavior and sympathetic control of cardiovascular function. For instance, it has been shown that PVN^MC4R+^→PBN pathway is responsible for feeding[23; 24; 39]; however, whether this pathway also affects cardiovascular function is unknown. Our anterograde tracing clearly shows that in addition to the PBN, PVN^MC4R+^ neurons also heavily innervate brain regions involved in sympathetic control of cardiovascular function, including NTS, LC, VML, and TSC. However, the functional roles of these projections in the sympathetic control of cardiovascular function have not yet been explored. Another important question that arose from present study is whether PVN^MC4R+^ neurons send collateral projections to different brain regions to simultaneously affect both feeding and autonomic control of cardiovascular function and thermoregulation. Much more refined functional circuit mapping studies are needed to answer these important yet unresolved questions in future.

There are some limitations to the present study. Although viral-mediated tract-tracing is widely used nowadays to delineate cell-type-specific brain circuits, whether most ChR2-eYFP fibers we observed make functional synaptic connections in those brain regions remains unclear. It is possible that some ChR2-eYFP fibers in certain brain regions could be just passing through axons. Comprehensive ChR2-assisted circuit mapping with electrophysiological recording in those projected brain regions may be required to prove the likelihood of true synaptic connections in the future. Additionally, it is extremely technically challenging to perfectly cover the entire rostral-to-caudal structure of PVN without contaminating adjacent brain regions by stereotaxic injection and therefore, neuroanatomical mapping of both efferent and afferent circuits we presented here could have been underestimated the actual input-output organization of PVN^MC4R+^ neurons. Additionally, although the number of mice used to evaluate BP and HR is admittedly low, we observed an explicit cardiovascular response upon DREADD activation of PVN^MC4R+^ neurons, strongly supporting the role of PVN^MC4R^ pathways in cardiovascular regulation. Another potential concern is the use of CNO as a mean for neuronal activation; it has been shown that when used in high dose (>5 mg/kg), CNO can be reverse-metabolized into clozapine[44; 45], an antipsychotic medication that could impact some behaviors and physiological measurements. While did not observe significant effects of CNO on feeding, BAT thermogenesis and cardiovascular parameters in control mice, it is worthy to note that tail temperature was slight but significantly increased by CNO treatment in control mice (although the effect was much smaller than that of DREADD activation of PVN^MC4R+^ neurons). Since the clozapine has been shown to have a vasodilatory effect[46; 47], we speculate that this small increase of tail temperature could be due to vasodilation caused by clozapine that converted from CNO. It is therefore worthwhile to test in future whether optogenetic activation of PVN^MC4R+^ neurons can replicate some of functional observations made in the present study.

## 5. CONCLUSION

In summary, we comprehensively mapped the neuroanatomical organization of PVN^MC4R+^ neurons and present further evidence to show that PVN^MC4R+^ circuits mediate much more complex behavioral and physiological regulations beyond feeding. PVN^MC4R^ signaling pathways have been the focus of appetite suppression for potential anti-obesity therapeutics. Understanding the neural basis of PVN^MC4R+^ signaling pathways for these complex behavioral and physiological regulations may therefore help to refine the strategy for targeting PVN^MC4R+^ to treat obesity and associated complications, such as obesity-associated hypertension.

## AUTHOR CONTRIBUTIONS

US and HC contributed to conceptualization, study design, methodology, and manuscript writing. US, KS, BAT, JED, JJ, SRR, YD, GD, BX, ZZ, and LVZ conducted and assisted the experiments. US and HC contributed to the data analysis.

## ACKNOWLEDGMENTS

We would like to thank Dr. Bradford Lowell (Beth Israel Deaconess Medical Center, Boston, MA) for the use of *MC4R*-Cre mouse line. This work was supported by grants from the National Institutes of Health (HL127673 and HL084207 to HC), the University of Iowa Fraternal Order of Eagles Diabetes Center Pilot Grant (to HC), and the American Heart Association (19POST34450083 to US, 19POST34380239 to GD).

## CONFLICT OF INTEREST

The authors declare no conflict of interest.

## Figures and legends

**Supplemental figure 1.**
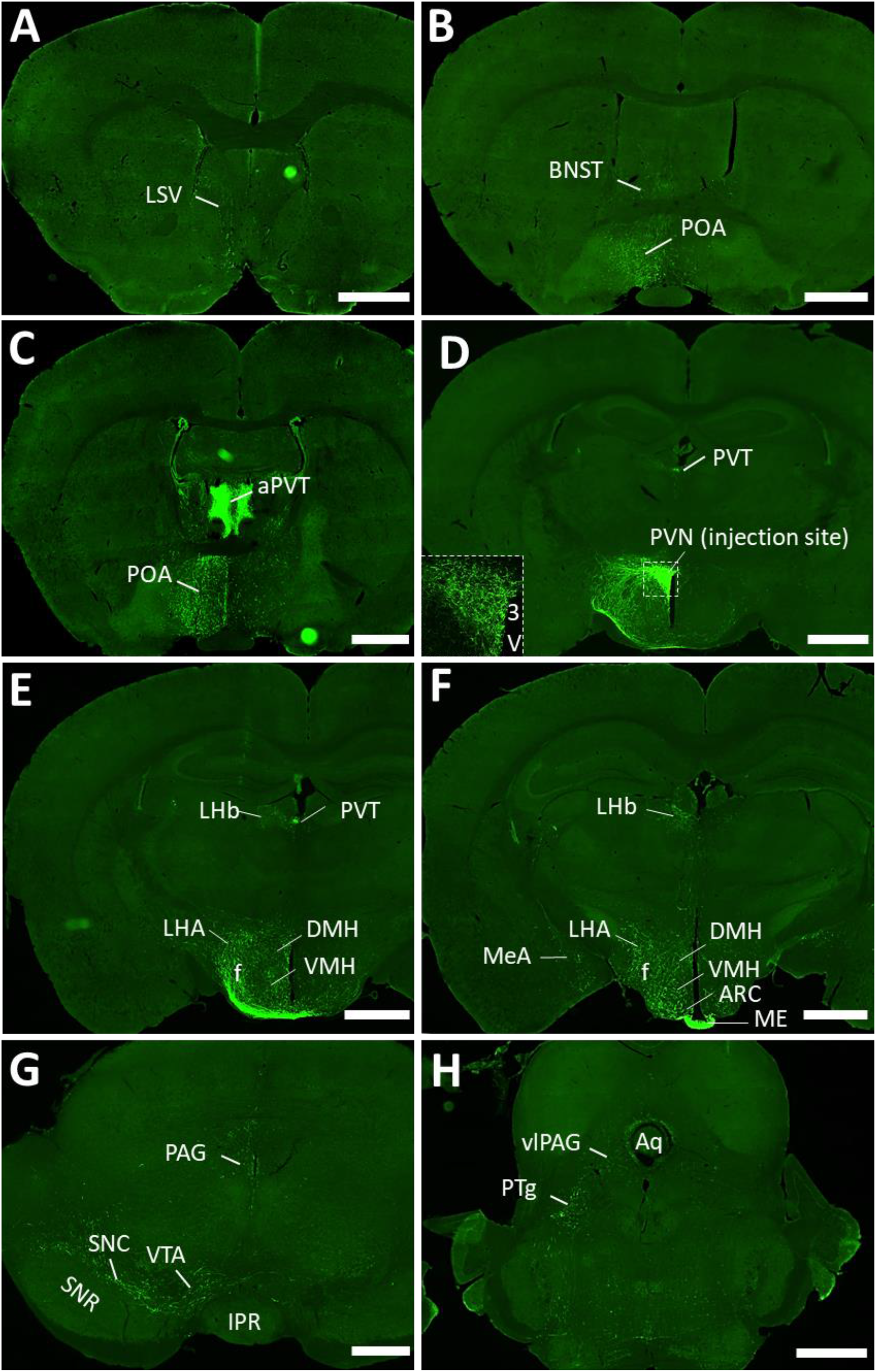
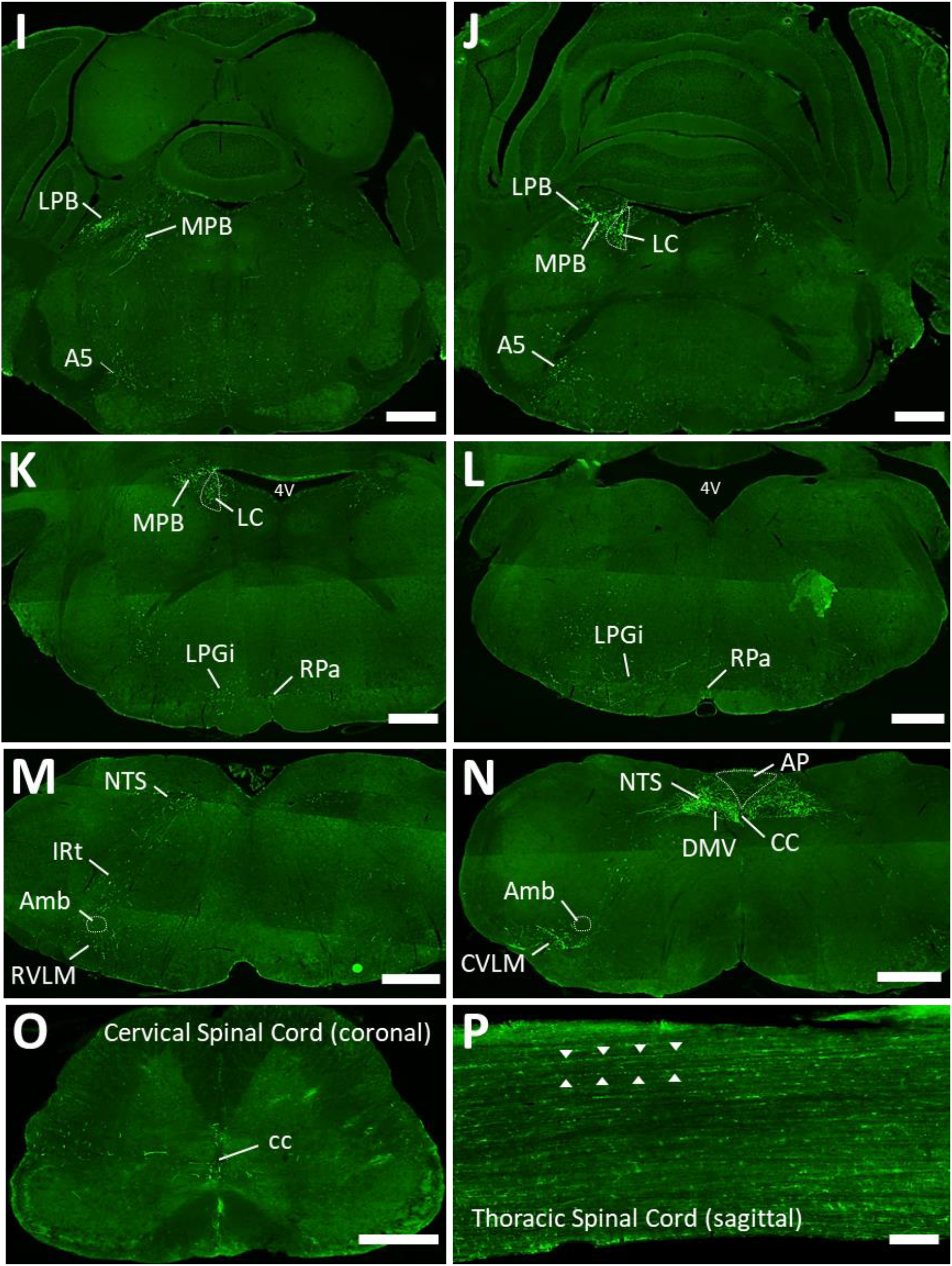
: Whole-brain mapping of efferent projections of PVNMC4R+ neurons. (A-N) Representative low magnification images of brain sections (from rostral to caudal order) showing eYFP projections to a variety of different brain regions. Note that the projections to most brain regions are predominantly unilateral, except aPVT, NTS and DMV where strong bilateral projections are observed. (O, P) Representative images of coronal section of upper cervical spinal cord (O) and sagittal section of thoracic spinal cord. White arrow heads indicate eYFP fibers traveling down the TSC (P). Abbreviations: bed nucleus of the stria terminalis (BNST), lateral (LS) and medial (MS) septal nuclei, preoptic area (POA), dorsomedial hypothalamic nucleus (DMH), ventromedial hypothalamic nucleus (VMH), lateral hypothalamic area (LHA), arcuate hypothalamic nucleus (ARC), median eminence (ME), medial amygdaloid nucleus (MeA), lateral habenular nucleus (LHb), paraventricular nucleus of the thalamus (PVT), posterior hypothalamic nucleus (PH), ventral premammillary nucleus (PMV), medial mammillary nucleus (MM), nucleus of the posterior commissure (Pcom), precommissural nucleus (Prc), periaqueductal grey (PAG), ventral tegmental area (VTA), substantia nigra (SN), subbrachial nucleus (SubB), dorsal raphe nucleus (DRN), ventrolateral PAG (vlPAG), parabrachial nucleus (PBN), locus coeruleus (LC), medial PBN (MBP), Gigantocellular reticular nucleus (Gi), ventrolateral medulla (VLM), nucleus of solitary tract (NTS), dorsal motor nucleus of the vagus (DMV), area postrema (AP), lateral reticular nucleus (LRt), thoracic spinal cord (TSC), lateral ventricle (LV), anterior commissure (aca), 3rd ventricle (3V), fornix (f), optic tract (opt), dorsal 3rd ventricle (D3V), (fr), mammillothalamic tract (mt), principal mammillary tract (pm), posterior commissure (pc), commissure of the superior colliculus (csc), aqueduct (Aq), 4th ventricle (4V), superior cerebellar peduncle (scp), perifacial zone (P7), ambiguus nucleus (Amb), central canal (CC). Scale bar: 1 mm for A-G and H, 500 µm for G and I-O, and 100 µm for P.

**Supplemental figure 2.**
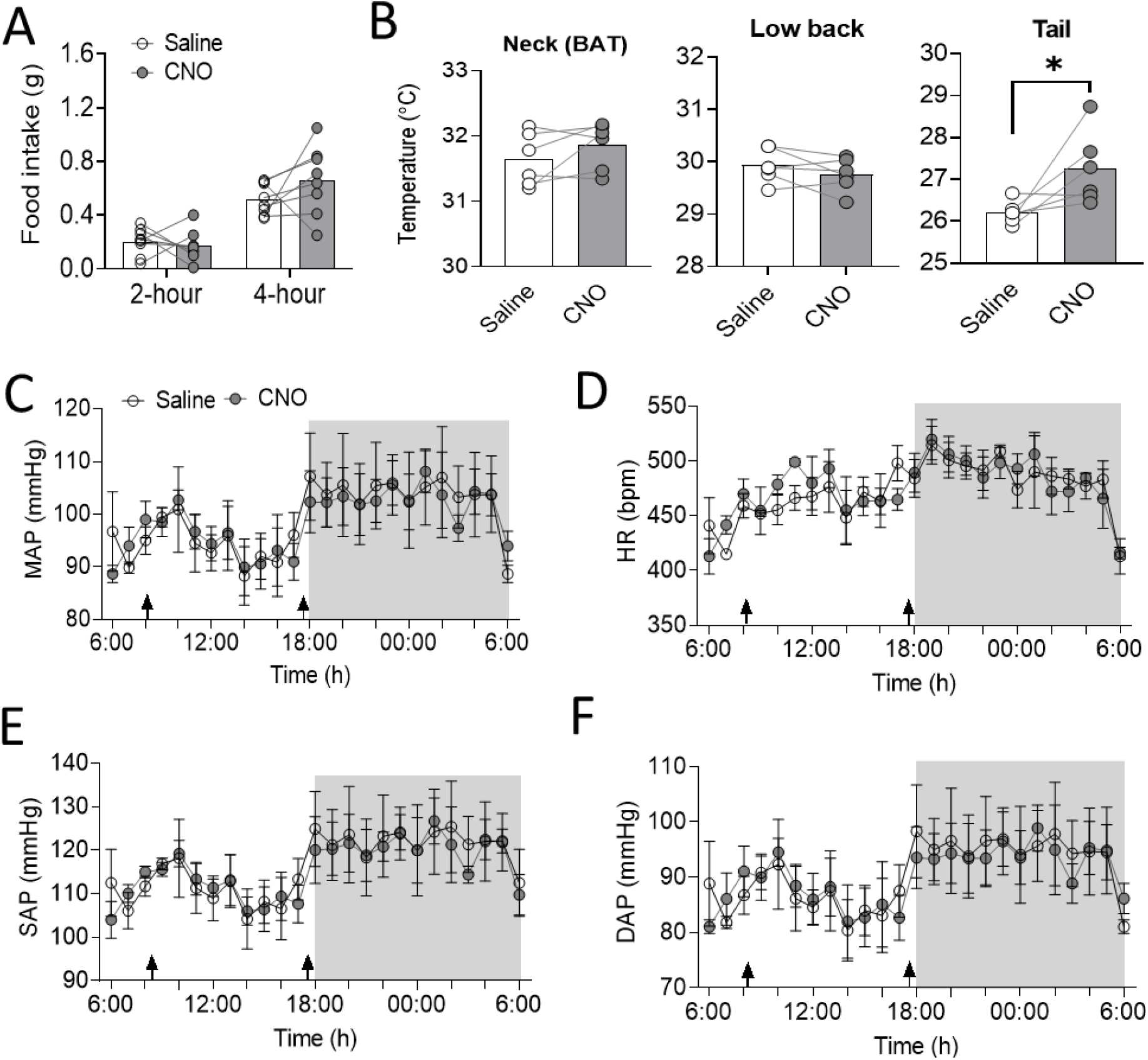
Evaluation of systemic CNO treatment (2 mg/kg) on feeding, thermoregulation and cardiovascular function in non-DREADD control mice. (A) Food intake following CNO injection right before the beginning of dark cycle (6 PM). (B) Body surface temperature measured by IR thermal imaging camera at baseline and after 1-hour of CNO treatment (2 mg/kg). (C-F) Chronic radio-telemetry recording of MAP (C), HR (D), SAP (E), and DAP (F) in response to saline and CNO treatment (n=3/group). *p<0.05 by paired Student’s t-test. Data are presented as mean ± SEM.

**Supplemental table 1:**
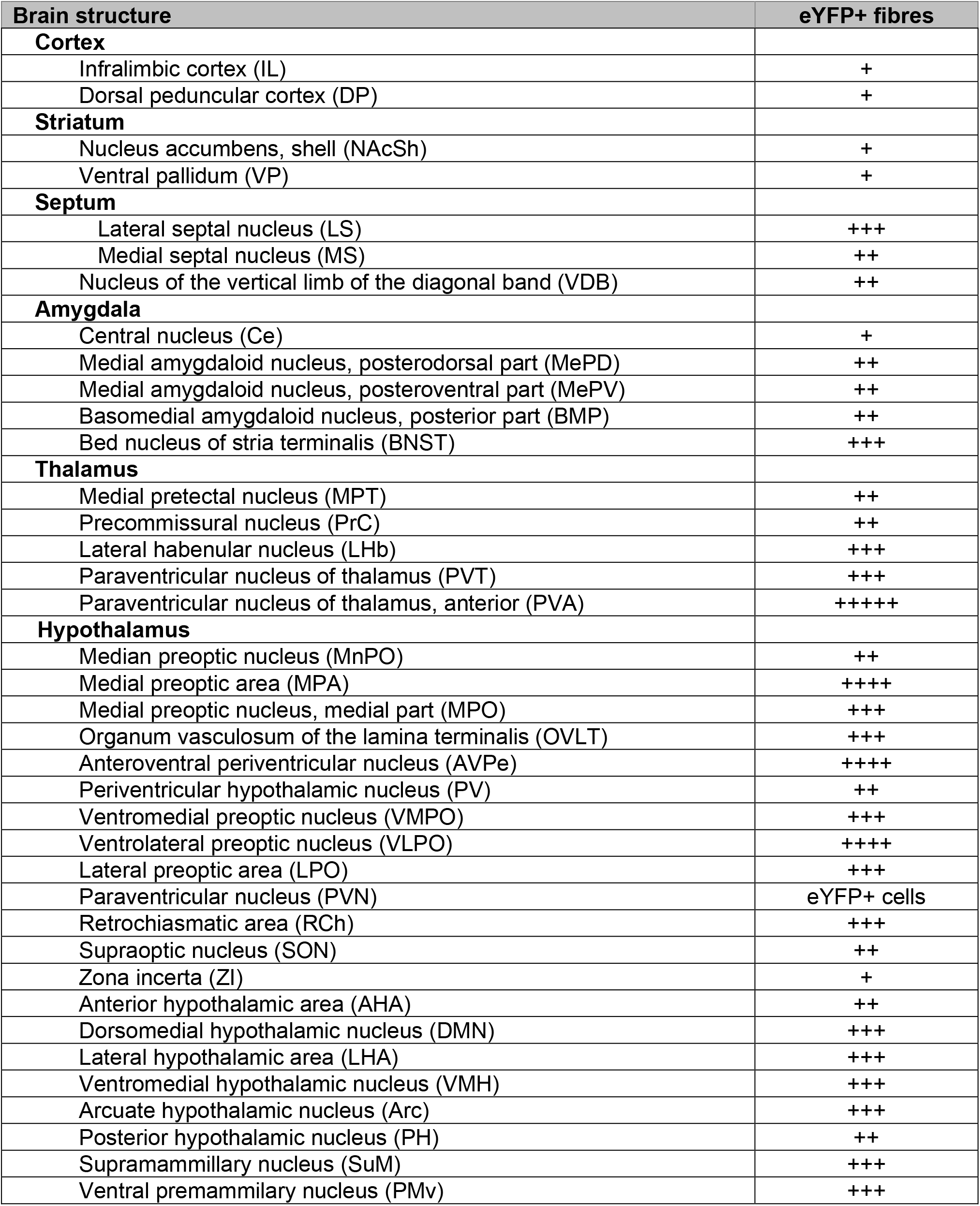
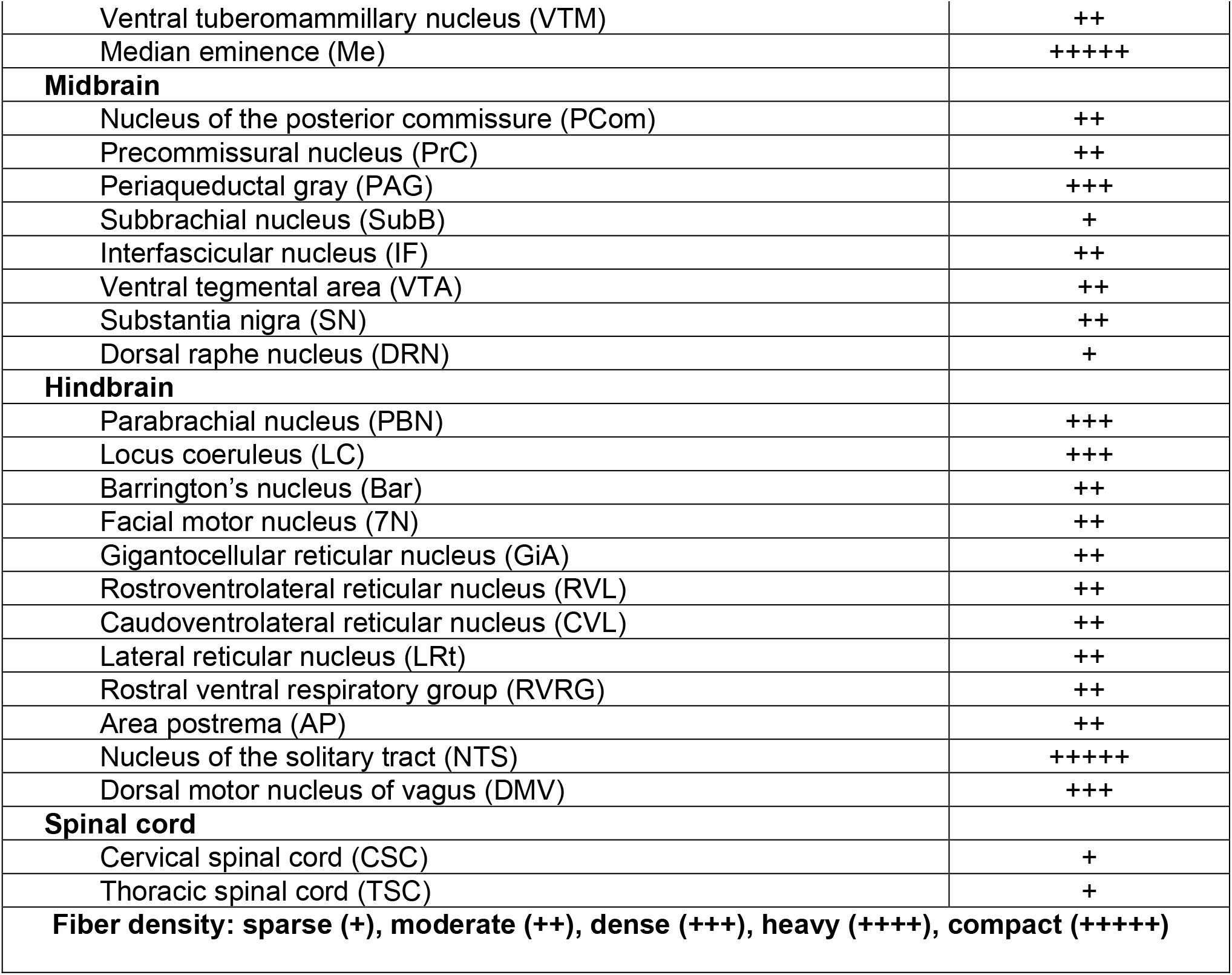

| **Brain structure** | **eYFP+ fibres** |
| --- | --- |
| **Cortex** |  |
| Infralimbic cortex (IL) | + |
| Dorsal peduncular cortex (DP) | + |
| **Striatum** |  |
| Nucleus accumbens, shell (NAcSh) | + |
| Ventral pallidum (VP) | + |
| **Septum** |  |
| Lateral septal nucleus (LS) | +++ |
| Medial septal nucleus (MS) | ++ |
| Nucleus of the vertical limb of the diagonal band (VDB) | ++ |
| **Amygdala** |  |
| Central nucleus (Ce) | + |
| Medial amygdaloid nucleus, posterodorsal part (MePD) | ++ |
| Medial amygdaloid nucleus, posteroventral part (MePV) | ++ |
| Basomedial amygdaloid nucleus, posterior part (BMP) | ++ |
| Bed nucleus of stria terminalis (BNST) | +++ |
| **Thalamus** |  |
| Medial pretectal nucleus (MPT) | ++ |
| Precommissural nucleus (PrC) | ++ |
| Lateral habenular nucleus (LHb) | +++ |
| Paraventricular nucleus of thalamus (PVT) | +++ |
| Paraventricular nucleus of thalamus, anterior (PVA) | +++++ |
| **Hypothalamus** |  |
| Median preoptic nucleus (MnPO) | ++ |
| Medial preoptic area (MPA) | ++++ |
| Medial preoptic nucleus, medial part (MPO) | +++ |
| Organum vasculosum of the lamina terminalis (OVLT) | +++ |
| Anteroventral periventricular nucleus (AVPe) | ++++ |
| Periventricular hypothalamic nucleus (PV) | ++ |
| Ventromedial preoptic nucleus (VMPO) | +++ |
| Ventrolateral preoptic nucleus (VLPO) | ++++ |
| Lateral preoptic area (LPO) | +++ |
| Paraventricular nucleus (PVN) | eYFP+ cells |
| Retrochiasmatic area (RCh) | +++ |
| Supraoptic nucleus (SON) | ++ |
| Zona incerta (ZI) | + |
| Anterior hypothalamic area (AHA) | ++ |
| Dorsomedial hypothalamic nucleus (DMN) | +++ |
| Lateral hypothalamic area (LHA) | +++ |
| Ventromedial hypothalamic nucleus (VMH) | +++ |
| Arcuate hypothalamic nucleus (Arc) | +++ |
| Posterior hypothalamic nucleus (PH) | ++ |
| Supramammillary nucleus (SuM) | +++ |
| Ventral premammillary nucleus (PMv) | +++ |
| Ventral tuberomammillary nucleus (VTM) | ++ |
| Median eminence (Me) | +++++ |
| **Midbrain** |  |
| Nucleus of the posterior commissure (PCom) | ++ |
| Precommissural nucleus (PrC) | ++ |
| Periaqueductal gray (PAG) | +++ |
| Subbrachial nucleus (SubB) | + |
| Interfascicular nucleus (IF) | ++ |
| Ventral tegmental area (VTA) | ++ |
| Substantia nigra (SN) | ++ |
| Dorsal raphe nucleus (DRN) | + |
| **Hindbrain** |  |
| Parabrachial nucleus (PBN) | +++ |
| Locus coeruleus (LC) | +++ |
| Barrington's nucleus (Bar) | ++ |
| Facial motor nucleus (7N) | ++ |
| Gigantocellular reticular nucleus (GiA) | ++ |
| Rostroventrolateral reticular nucleus (RVL) | ++ |
| Caudoventrolateral reticular nucleus (CVL) | ++ |
| Lateral reticular nucleus (LRt) | ++ |
| Rostral ventral respiratory group (RVRG) | ++ |
| Area postrema (AP) | ++ |
| Nucleus of the solitary tract (NTS) | +++++ |
| Dorsal motor nucleus of vagus (DMV) | +++ |
| **Spinal cord** |  |
| Cervical spinal cord (CSC) | + |
| Thoracic spinal cord (TSC) | + |
| **Fiber density: sparse (+), moderate (++), dense (+++), heavy (++++), compact (+++++)** |  |

Abbreviations:

## Abbreviations

AP: Area postrema
ARC: Arcuate nucleus
BNST: Bed nucleus of the stria terminalis BP Blood pressure
ChR2: Channelrhodopsin-2
CNO: Clozapine-N-oxide
CVLM: Caudal ventrolateral medulla DAP Diastolic arterial pressure
DMH: Dorsomedial nucleus of hypothalamus DMV Dorsal motor nucleus of the vagus
DREADD: Designer Receptors Exclusively Activated by Designer Drugs Flp Flippase
HR: Heart rate
IHC: Immunohistochemistry
LC: Locus coeruleus
LHA: Lateral hypothalamic area
MAP: Mean arterial pressure
MC4R: Melanocortin 4 receptor
ME: Median eminence
NTS: Nucleus of solitary tract
PAG: Periaqueductal gray
PBN: Parabrachial nucleus
POA: Preoptic area
PVN: Paraventricular nucleus of hypothalamus PVT Paraventricular nucleus of thalamus
RV: Rabies virus
RV-EnvA-ΔG: EnvA-pseudotyped G-deleted Rabies viruses RVLM Rostral ventrolateral medulla
SAP: Systolic arterial pressure
TSC: Thoracic spinal cord
VMH: Ventromedial nucleus of hypothalamus vSUB Ventral subiculum

## Notes

### Competing Interest Statement

The authors have declared no competing interest.

## REFERENCES

[1] Krauss, R.M., Winston, M., Fletcher, B.J., Grundy, S.M., 1998. Obesity: impact on cardiovascular disease. Circulation 98(14):1472–1476.

[2] Tentolouris, N., Liatis, S., Katsilambros, N., 2006. Sympathetic system activity in obesity and metabolic syndrome. Ann N Y Acad Sci 1083:129–152.

[3] Grassi, G., Biffi, A., Seravalle, G., Trevano, F.Q., Dell’Oro, R., Corrao, G., et al., 2019. Sympathetic Neural Overdrive in the Obese and Overweight State. Hypertension 74(2):349–358.

[4] do Carmo, J.M., da Silva, A.A., Wang, Z., Fang, T., Aberdein, N., Perez de Lara, C.E., et al., 2017. Role of the brain melanocortins in blood pressure regulation. Biochim Biophys Acta Mol Basis Dis 1863(10 Pt A):2508–2514.

[5] da Silva, A.A., do Carmo, J.M., Hall, J.E., 2013. Role of leptin and central nervous system melanocortins in obesity hypertension. Curr Opin Nephrol Hypertens 22(2):135–140.

[6] da Silva, A.A., do Carmo, J.M., Wang, Z., Hall, J.E., 2019. Melanocortin-4 Receptors and Sympathetic Nervous System Activation in Hypertension. Curr Hypertens Rep 21(6):46.

[7] da Silva, A.A., do Carmo, J.M., Wang, Z., Hall, J.E., 2014. The brain melanocortin system, sympathetic control, and obesity hypertension. Physiology (Bethesda) 29(3):196–202.

[8] Huszar, D., Lynch, C.A., Fairchild-Huntress, V., Dunmore, J.H., Fang, Q., Berkemeier, L.R., et al., 1997. Targeted disruption of the melanocortin-4 receptor results in obesity in mice. Cell 88(1):131–141.

[9] Yeo, G.S., Farooqi, I.S., Aminian, S., Halsall, D.J., Stanhope, R.G., O’Rahilly, S., 1998. A frameshift mutation in MC4R associated with dominantly inherited human obesity. Nat Genet 20(2):111–112.

[10] Kuo, J.J., Silva, A.A., Hall, J.E., 2003. Hypothalamic melanocortin receptors and chronic regulation of arterial pressure and renal function. Hypertension 41(3 Pt 2):768–774.

[11] Rahmouni, K., Haynes, W.G., Morgan, D.A., Mark, A.L., 2003. Role of melanocortin-4 receptors in mediating renal sympathoactivation to leptin and insulin. J Neurosci 23(14):5998–6004.

[12] Tallam, L.S., da Silva, A.A., Hall, J.E., 2006. Melanocortin-4 receptor mediates chronic cardiovascular and metabolic actions of leptin. Hypertension 48(1):58–64.

[13] Greenfield, J.R., Miller, J.W., Keogh, J.M., Henning, E., Satterwhite, J.H., Cameron, G.S., et al., 2009. Modulation of blood pressure by central melanocortinergic pathways. N Engl J Med 360(1):44–52.

[14] Greenfield, J.R., 2011. Melanocortin signalling and the regulation of blood pressure in human obesity. J Neuroendocrinol 23(2):186–193.

[15] Tallam, L.S., Stec, D.E., Willis, M.A., da Silva, A.A., Hall, J.E., 2005. Melanocortin-4 receptor-deficient mice are not hypertensive or salt-sensitive despite obesity, hyperinsulinemia, and hyperleptinemia. Hypertension 46(2):326–332.

[16] Sayk, F., Heutling, D., Dodt, C., Iwen, K.A., Wellhoner, J.P., Scherag, S., et al., 2010. Sympathetic function in human carriers of melanocortin-4 receptor gene mutations. J Clin Endocrinol Metab 95(4):1998–2002.

[17] do Carmo, J.M., da Silva, A.A., Hall, J.E., 2015. Role of hindbrain melanocortin-4 receptor activity in controlling cardiovascular and metabolic functions in spontaneously hypertensive rats. J Hypertens 33(6):1201–1206.

[18] Haynes, W.G., Morgan, D.A., Djalali, A., Sivitz, W.I., Mark, A.L., 1999. Interactions between the melanocortin system and leptin in control of sympathetic nerve traffic. Hypertension 33(1 Pt 2):542–547.

[19] Qin, C., Li, J., Tang, K., 2018. The Paraventricular Nucleus of the Hypothalamus: Development, Function, and Human Diseases. Endocrinology 159(9):3458–3472.

[20] Kishi, T., Aschkenasi, C.J., Lee, C.E., Mountjoy, K.G., Saper, C.B., Elmquist, J.K., 2003. Expression of melanocortin 4 receptor mRNA in the central nervous system of the rat. J Comp Neurol 457(3):213–235.

[21] Liu, H., Kishi, T., Roseberry, A.G., Cai, X., Lee, C.E., Montez, J.M., et al., 2003. Transgenic mice expressing green fluorescent protein under the control of the melanocortin-4 receptor promoter. J Neurosci 23(18):7143–7154.

[22] Balthasar, N., Dalgaard, L.T., Lee, C.E., Yu, J., Funahashi, H., Williams, T., et al., 2005. Divergence of melanocortin pathways in the control of food intake and energy expenditure. Cell 123(3):493–505.

[23] Shah, B.P., Vong, L., Olson, D.P., Koda, S., Krashes, M.J., Ye, C., et al., 2014. MC4R-expressing glutamatergic neurons in the paraventricular hypothalamus regulate feeding and are synaptically connected to the parabrachial nucleus. Proc Natl Acad Sci U S A 111(36):13193–13198.

[24] Garfield, A.S., Li, C., Madara, J.C., Shah, B.P., Webber, E., Steger, J.S., et al., 2015. A neural basis for melanocortin-4 receptor-regulated appetite. Nat Neurosci 18(6):863–871.

[25] Li, P., Cui, B.P., Zhang, L.L., Sun, H.J., Liu, T.Y., Zhu, G.Q., 2013. Melanocortin 3/4 receptors in paraventricular nucleus modulate sympathetic outflow and blood pressure. Exp Physiol 98(2):435–443.

[26] Li, Y.Q., Shrestha, Y., Pandey, M., Chen, M., Kablan, A., Gavrilova, O., et al., 2016. G(q/11)alpha and G(s)alpha mediate distinct physiological responses to central melanocortins. J Clin Invest 126(1):40–49.

[27] Ye, Z.Y., Li, D.P., 2011. Activation of the melanocortin-4 receptor causes enhanced excitation in presympathetic paraventricular neurons in obese Zucker rats. Regul Pept 166(1-3):112–120.

[28] Cui, H., Sohn, J.W., Gautron, L., Funahashi, H., Williams, K.W., Elmquist, J.K., et al., 2012. Neuroanatomy of melanocortin-4 receptor pathway in the lateral hypothalamic area. J Comp Neurol 520(18):4168–4183.

[29] Vialou, V., Cui, H., Perello, M., Mahgoub, M., Yu, H.G., Rush, A.J., et al., 2011. A role for DeltaFosB in calorie restriction-induced metabolic changes. Biol Psychiatry 70(2):204–207.

[30] Morgan, D.A., McDaniel, L.N., Yin, T., Khan, M., Jiang, J., Acevedo, M.R., et al., 2015. Regulation of glucose tolerance and sympathetic activity by MC4R signaling in the lateral hypothalamus. Diabetes 64(6):1976–1987.

[31] Cui, H., Lu, Y., Khan, M.Z., Anderson, R.M., McDaniel, L., Wilson, H.E., et al., 2015. Behavioral disturbances in estrogen-related receptor alpha-null mice. Cell Rep 11(3):344–350.

[32] Zhu, Z., Sierra, A., Burnett, C.M., Chen, B., Subbotina, E., Koganti, S.R., et al., 2014. Sarcolemmal ATP-sensitive potassium channels modulate skeletal muscle function under low-intensity workloads. J Gen Physiol 143(1):119–134.

[33] Zingman, L.V., Hodgson, D.M., Bast, P.H., Kane, G.C., Perez-Terzic, C., Gumina, R.J., et al., 2002. Kir6.2 is required for adaptation to stress. Proc Natl Acad Sci U S A 99(20):13278–13283.

[34] Jiang, J., Morgan, D.A., Cui, H., Rahmouni, K., 2020. Activation of hypothalamic AgRP and POMC neurons evokes disparate sympathetic and cardiovascular responses. Am J Physiol Heart Circ Physiol 319(5):H1069–H1077.

[35] Pham, J., Cabrera, S.M., Sanchis-Segura, C., Wood, M.A., 2009. Automated scoring of fear-related behavior using EthoVision software. J Neurosci Methods 178(2):323–326.

[36] Deacon, R.M., 2006. Digging and marble burying in mice: simple methods for in vivo identification of biological impacts. Nat Protoc 1(1):122–124.

[37] Krashes, M.J., Shah, B.P., Madara, J.C., Olson, D.P., Strochlic, D.E., Garfield, A.S., et al., 2014. An excitatory paraventricular nucleus to AgRP neuron circuit that drives hunger. Nature 507(7491):238–242.

[38] Li, C., Navarrete, J., Liang-Guallpa, J., Lu, C., Funderburk, S.C., Chang, R.B., et al., 2019. Defined Paraventricular Hypothalamic Populations Exhibit Differential Responses to Food Contingent on Caloric State. Cell Metab 29(3):681–694 e685.

[39] Li, M.M., Madara, J.C., Steger, J.S., Krashes, M.J., Balthasar, N., Campbell, J.N., et al., 2019. The Paraventricular Hypothalamus Regulates Satiety and Prevents Obesity via Two Genetically Distinct Circuits. Neuron 102(3):653–667 e656.

[40] Song, C.K., Vaughan, C.H., Keen-Rhinehart, E., Harris, R.B., Richard, D., Bartness, T.J., 2008. Melanocortin-4 receptor mRNA expressed in sympathetic outflow neurons to brown adipose tissue: neuroanatomical and functional evidence. Am J Physiol Regul Integr Comp Physiol 295(2):R417–428.

[41] Voss-Andreae, A., Murphy, J.G., Ellacott, K.L., Stuart, R.C., Nillni, E.A., Cone, R.D., et al., 2007. Role of the central melanocortin circuitry in adaptive thermogenesis of brown adipose tissue. Endocrinology 148(4):1550–1560.

[42] Xu, Y., Wu, Z., Sun, H., Zhu, Y., Kim, E.R., Lowell, B.B., et al., 2013. Glutamate mediates the function of melanocortin receptor 4 on Sim1 neurons in body weight regulation. Cell Metab 18(6):860–870.

[43] Tsuchiya, K., 2001. The Rat Tail as a Model Organ for Peripheral Vasodilation.192–199.

[44] Manvich, D.F., Webster, K.A., Foster, S.L., Farrell, M.S., Ritchie, J.C., Porter, J.H., et al., 2018. The DREADD agonist clozapine N-oxide (CNO) is reverse-metabolized to clozapine and produces clozapine-like interoceptive stimulus effects in rats and mice. Sci Rep 8(1):3840.

[45] Gomez, J.L., Bonaventura, J., Lesniak, W., Mathews, W.B., Sysa-Shah, P., Rodriguez, L.A., et al., 2017. Chemogenetics revealed: DREADD occupancy and activation via converted clozapine. Science 357(6350):503–507.

[46] Mateus, L.S., Albuquerque, A.A.S., Celotto, A.C., Evora, P.R.B., 2019. In vitro evidence that endothelium-dependent vasodilatation induced by clozapine is mediated by an ATP-sensitive potassium channel. Pharmacol Rep 71(3):522–527.

[47] Boussery, K., Lambrecht, S., Delaey, C., Van de Voorde, J., 2005. Clozapine directly relaxes bovine retinal arteries. Curr Eye Res 30(2):139–146.

